# Lectins in pistils play a key role in self-incompatibility in the heterostylous *Linum perenne*

**DOI:** 10.1101/2023.04.02.535286

**Authors:** Hanna Levchuk, Yuliia Makhno, Maria Manuela Ribeiro Costa, Michael Lenhard

## Abstract

Self-incompatibility is one of the mechanisms preventing inbreeding in populations, based on the failure of pollen grains to germinate or of pollen tubes to grow normally in incompatible crosses. A unique model for studying this process are heterostylous species, in which self-incompatibility alleles are closely linked to flower morphological traits. One such species is *L. perenne*, a classic distylous species with reciprocal herkogamy and complete self-incompatibility strongly linked. The mechanism of self-pollen recognition is poorly understood in most heterostylous species, including *L. perenne*. As self-incompatibility in other systems is based on protein-protein interactions and many extracellular proteins are glycosylated, one of the possible participants in this process may be lectins. Therefore, we aimed to investigate the lectin activity in pistils and stamens of *L. perenne* L-morph and S-morph plants and to establish whether these lectins take part in self-incompatibility. We show that pistil-derived galactose-specific Ca^2+^-independent lectins from the respective other morph can overcome self-incompatibility in heterostylous *L. perenne*. This may be due to the presence of some lectins that are specific for arabinose and xylose. Our results suggest that lectins participate in signaling pathways for recognition at the pollen-stigma interface and are involved in the regulation of the self-incompatibility process in *L. perenne*.

## Introduction

Self-incompatibility is a major mechanism to prevent inbreeding, based on the specific discrimination between self and non-self-pollen (Iwano and Takayama, 2012). Since inbreeding can lead to inbreeding depression and reduced potential for evolutionary adaptation, and eventually result in the extinction of species, the effectiveness of self-incompatibility (SI) systems is an important contributor to the evolutionary success of flowering plants (Lewis, 1979; Harris et al., 1984; Cornish et al., 1988). Self-incompatible crosses are characterized by the failure of pollen germination or the arrest of pollen tube growth (Schopfer et al., 1999). The genetic control of this process is based on a single chromosomal locus with several different *S* haplotypes that are responsible for protein-protein recognition between male and female partners.

There are three main types of SI systems: homomorphic gametophytic, homomorphic sporophytic and heteromorphic self-incompatibility. Gametophytic incompatibility is genetically controlled by the haploid (hence gametophytic) genome of the pollen grain and the diploid genome of the pistil tissue (Takasaki et al., 2000, Iwano and Takayama, 2012). By contrast, in sporophytic self-incompatibility the interaction between pollen and pistil is genetically controlled by the diploid (hence sporophyte) genome of the parent plant in which the pollen developed and the diploid genotype of the pistil tissue (McClure et al., 1989; Boskovic and Tobutt, 1996; Xue et al., 1996). The gametophytic SI mechanism in Solanaceae and the sporophytic SI systems in Brassicaceae and Papaveraceae are understood in great molecular detail by now (Fujii et al., 2016).

In contrast to the above homomorphic SI systems where the different mating types are not morphologically differentiated, in heteromorphic incompatibility the *S* loci are also associated with flower morphological traits. As a result, in populations of heterostylous species there are two (sometimes three) flower morphs that differ in the length of the style and stamen filaments and represent different mating types. The morphology of the flowers with reciprocal herkogamy represents a topological barrier that promotes efficient cross-pollination between the two morphs and thus prevents pollen wastage (Lloyd and Webb, 1992). In most heterostylous species, a physiological self-incompatibility barrier prevents seed set after self- and intra-morph pollination. Heteromorphic self-incompatibility occurs in at least 24 families and more than 164 genera (Ganders, 1979). The biology and genetics are described for several species; for example, *Linum* (Lewis, 1943; Ghosh and Shivanna, 1980) and *Primula* (Heslop-Harrison and Shivanna, 1981; Shivanna et al., 1981), but the molecular mechanism and its genetic control remain poorly understood.

Several researchers have attempted to elucidate the mechanisms of incompatibility including Lewis (1943), Dulberger (1974; 1975a, b; 1987), Golynskaya et al. (1976), Ghosh and Shivanna (1980), Shivanna et al. (1981; 1983), Stevens and Murray (1982), Schou and Ban (1985) and Murray (1986) and reviewed by Dulberger (1992). More recently, Wong et al. (1994) discovered a style-specific protein apparently unique to long-styled plants of *Auerrhoa carambola* (Oxalidaceae). In some distylous species of genus *Turnera* morph-specific proteins were found in pollen and styles (Athanasiou et al., 1997). Also, morph-specific differential expression of certain genes in pistils and stamens of some heterostylous species has been reported, in particular in buckwheat (Yasui et al., 2012), *Linum grandiflorum* (Ushijima et al., 2012) and *Primula* species (Nowak et al., 2015; Li et al., 2011.) Genetic studies in *Primula* have recently shown an important role for brassinosteroid levels in determining the female incompatibility type, yet the molecular mechanisms translating different brassinosteroid levels into different compatibility responses remain to be elucidated (Huu et al., 2022). Similarly, the *S*-locus specific auxin-biosynthesis gene *YUCCA6* in *Turnera* has been shown to be important for determining male incompatibility type (Henning, Shore and McCubbin, 2022). However, despite this progress, no clear picture of the mechanism(s) of incompatibility has yet emerged for distylous species.

As the basis of the self-incompatibility mechanism is protein-protein interaction (Wong et al., 1994, Ushijima et al., 2012), and many extracellular proteins are glycosylated, one of the possible participants in this process can be lectins. This is a group of glycoproteins that are capable of selectively and reversibly binding carbohydrates on cell surfaces (Peumans and Van Damme, 1995, Sharon and Lis, 2004). Based on the analysis of protein sequences and of genome and transcriptome data (Dang and Van Damme, 2016, Eggermont et al., 2017), lectin domains were identified (Van Damme et al., 1998, Tsaneva and Van Damme, 2020), which are evolutionarily conserved regions that may contain certain motifs associated with the carbohydrate binding site (Eggermont et al., 2017). Based on the presence of a typical lectin domain (Van Damme et al., 1998, Tsaneva and Van Damme, 2020, De Coninck and Van Damme, 2021.), all plant lectins have been grouped into 12 families, which are characterized by their molecular structure and carbohydrate binding properties (Van Damme et al., 2008). In addition to carbohydrate-binding domains, proteins of this group are also capable of binding proteins and lipids (De Coninck and Van Damme, 2021). Due to their carbohydrate binding ability, lectins play an important role in the control of intracellular transport processes and intercellular recognition (Rudiger and Gabius, 2001), including pollination. Therefore, lectins can be one of the keys in the regulation of self-incompatibility. Indeed, in some of the earlier studies (Golynskaya et al., 1976; Shivanna et al.,1981, 1983), pistil and stamen lectins were suggested to be involved in regulating self-incompatibility and the pollination process. More recently, it was found that lectins of some species are directly involved in the growth of the pollen tube *in vitro* (Matveeva et al., 2007; Sousa et al., 2013). Supporting a role for lectins and carbohydrate binding in the self-incompatibility response, the female determinant of SI in the Brassicaceae, the *S*-locus receptor kinase SRK contains two lectin GNA [*Galanthus nivalis* (snowdrop) agglutinin] domains at its N-terminal end (Ma et al., 2016; Bellande et al., 2017). The genus *Linum* provides an attractive system to study pollination mechanisms and heteromorphic SI, as it contains both homostylous and heterostyled species with different types of self-incompatibility. Several studies have been carried out on the breeding system of *Linum* species (Ockendon, 1968; Dulberger, 1973, 1981; Rogers, 1979). Darwin (1877) described the existence of distyly in several species such as *L. pubescens* Banks & Sol., *L. grandiflorum* Desf., *L. mucronatum* Bertol., *L. flavum* L., *L. perenne* L., *L. austriacum* L., and *L. maritimum* L. Of these, the European blue flax *Linum perenne* is a classic distylous species with reciprocal herkogamy and complete self-incompatibility (Darwin, 1864). For this reason, it was chosen as the object of the present study. We aimed to investigate the lectin activity in pistils and stamens of *L. perenne* L-morphs and S-morphs and to establish whether these lectins take part in self-incompatibility. The main purpose was to establish whether self-incompatibility can be overcome by treating stigmas before pollination with the lectins from the compatible floral morph.

## Materials and methods

### Plant material

Flax plants (*Linum perenne* L.) were grown under standard greenhouse conditions in the Portuguese Bank of Plant Germplasm (Braga, Portugal) under natural light and temperature conditions. For extracting lectins, both pistils and stamens of unpollinated flowers were collected immediately after the opening of the flowers (during two hours after the opening of the flower), frozen in liquid nitrogen and stored at −70^0^C. Pistils and stamens of different floral morphs (from short-styled (S-morph) and long-styled (L-morph) flowers) were individually collected (Fig. 1).

**Fig. 1.**
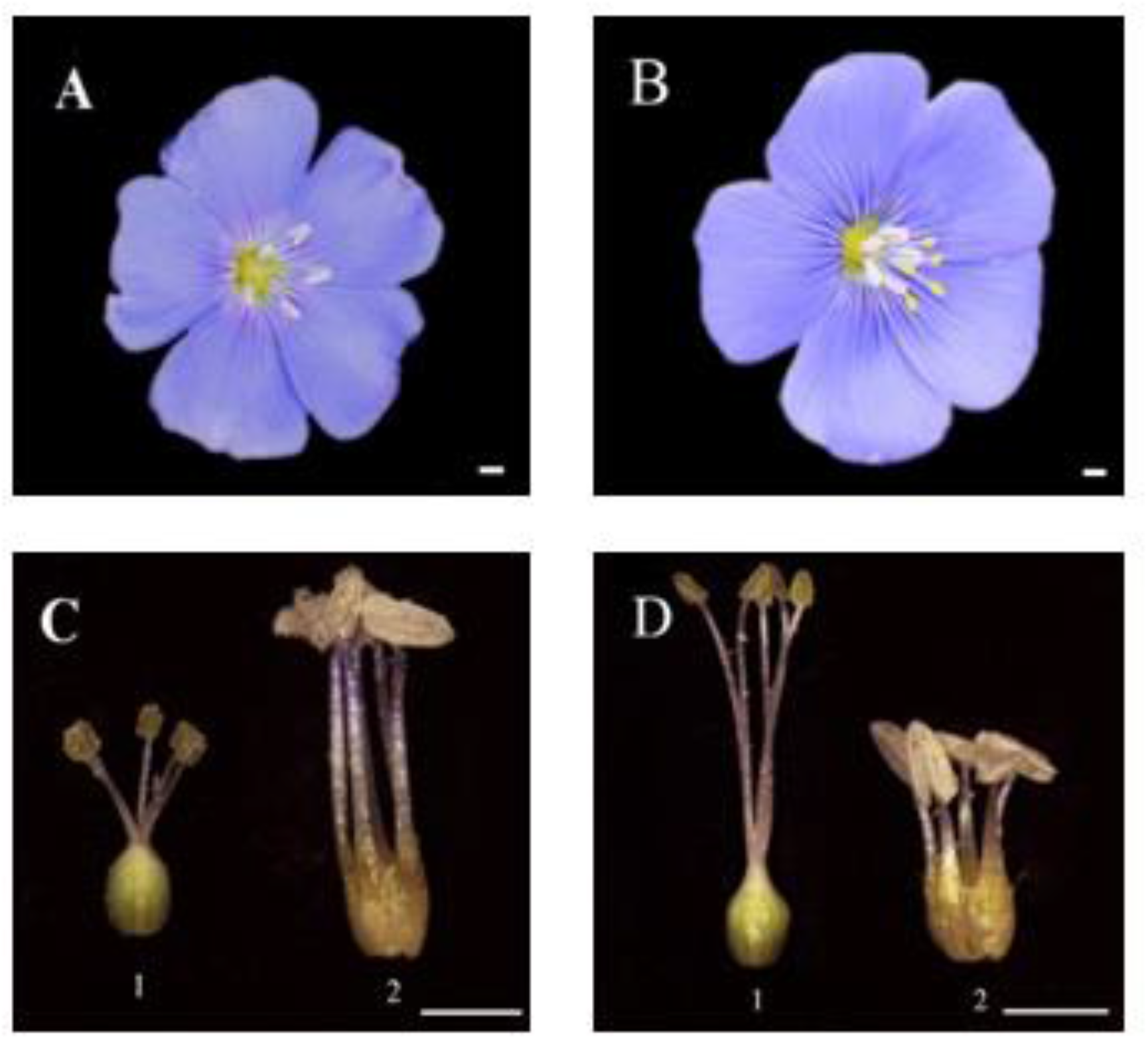
Flowers of the different morphs of the heterostyled species *L. perenn*e. (A) *L. perenne* short-styled (S) floral morph; (B) *L. perenne* long-styled (L) floral morph; (C and D) pistils (1) and stamens (2) of S-morphs (C) and L-morphs (D) of *L. perenne*. Scale bars in all pictures are 2 mm.

### Lectin extracts and purification

The lectins were extracted separately from the pistils and stamens of L-morphs and S-morphs of *L. perenne* (Fig. 1). Lectins from different cell fractions were obtained: cytoplasm (soluble lectins), membranes (membrane lectins) and cell walls (cell wall lectins). For purification of lectins from other proteins and the separation of individual lectins according to their carbohydrate specificity, the initial protein extracts were purified by lectin binding to glucose-containing and galactose-containing resins. Lectins were eluted from resins using carbohydrate solutions – glucose or galactose respectively (Ca^2+^-independent lectin) – and EDTA (Ca^2+^-dependent lectins).

Lectins were extracted from various cell fractions (membranes, soluble cytoplasm, and cell walls) according to standard protocols (Levchuk et al., 2014): cellular fractions were separated by centrifugation as described below, and lectins were isolated with 0.02 M potassium phosphate buffer at pH 6.8 with the addition of 0.35 M sucrose, 0.1 M ascorbic acid, and 0.01 M EDTA. A sample (0.15 g) was homogenized with 2 mL of 20 mM potassium phosphate buffer (pH 6.8) and left for extraction for 20 min at 4 °С with shaking, and then centrifuged for 15 min at 10 000 g. Supernatant (the fraction enriched in soluble lectins) was used for subsequent analysis. In order to isolate membranes from the organelle pellet containing nuclei, mitochondria, and plastids, it was homogenized with 0.2 mL of initial potassium phosphate buffer (pH 6.8) with the addition of Triton X-100 to 0.05% and left for extraction for 2 h at 4°С with shaking. The suspension was centrifuged for 15 min at 10 000 g; the resulting supernatant contained membrane lectins and the pellet contained cell walls. In order to separate cell walls, the cell wall pellet was homogenized with 2 mL of 20 mM potassium phosphate buffer (pH 4.0) supplemented with Triton X-100 detergent to 0.05%, left for extraction for 12 h at 4°С with shaking. The suspension was centrifuged similarly; the resulting supernatant contained cell wall lectins. Solubilized lectins of all the fractions were concentrated with 70% ammonium sulphate, and the pellets were dissolved in 100 µL of PBS and dialyzed using centrifugal filters (Amicon Ultra-15 Centrifugal Filter Units (30 000 NMWL)).

Obtained lectin extracts were purified and separated into several fractions by affinity chromatography sequentially using a glucose-containing resin (Sephacryl S100) and a galactose-containing resin (Superose 6). Prior to sample injection the resins were washed using a 10-fold volume of PBS. Each lectin extract (100 µL) was incubated with 500 µL of Sephacryl S100 for 2 h at 4°С with shaking. After centrifugation (5 min at 1 000 g and 4 ^°^C) the supernatant was incubated with 500 µL of Superose 6 for 2 h at 4°С with shaking centrifugation (5 min at 1 000 g and 4^°^C). From the Sephacryl and superpose resins, the glucose-binding and galactose-binding Ca^2+^-independent lectins were eluted initially using 10% carbohydrate (glucose or galactose respectively) and then (after washing the resin) using 0.01 M EDTA to elute the Ca^2+^-dependent lectins from these resins. The eluates were dialyzed against PBS with three changes and concentrated to 100 µL using centrifugal filters. For the pistil treatments the lectin extracts were diluted to titer 1:4. The eluted fractions and lectin extracts on each step of purification were monitored for lectin activity and protein content.

### Determination of lectin activity

Lectin extracts were characterized by quantitative (lectin activity) and qualitative (carbohydrate specificity) indices. Lectin activity was determined by the hemagglutination assay with a 2% suspension of rabbit erythrocytes. Hemagglutination tests were carried out in microtiter 96-well (U-shaped) plates. The lectin was diluted serially (2-fold), adjusting the sample volume in each well to 50 µL with PBS. Diluted samples were each mixed with 50 µL of the 2% suspension of rabbit erythrocytes. The reaction mixture was incubated for 1 h at room temperature and then observed for agglutination. Protein concentration was determined according to Lowry et al. (1951) using bovine serum albumin as a standard. The total hemagglutination activity was determined as hemagglutination unit (U) or titer and was defined as the reciprocal of the highest dilution showing detectable agglutination. The specific lectin activity was determined as the U per mg of protein. To determine the sugar-binding specificity of the lectin, 10 different sugars including pentoses, hexoses, disaccharides and amino-sugars were tested for their ability to inhibit lectin induced hemagglutination. For this the lectin was diluted serially to titer 1:4 and in each well 50 µL of this lectin solution and 50 µL of 0.6 M carbohydrate solution were mixed. After 20 min 50 µL of a 2% suspension of rabbit erythrocytes was added to each well. The reaction mixture was incubated for 1 h at room temperature and then observed for inhibition of agglutination in the presence of certain carbohydrates.

### Genetic crosses

Crosses on recently opened flowers were made with fresh pollen collected from plants of known floral morphs. Crosses were classified as fully incompatible if there was no capsule formation after pollination. Pollen tube growth was monitored by staining the styles with aniline blue about 2 h after pollination, followed by observations on a fluorescence microscope. To establish the ability of lectins to overcome the self-incompatibility during self-pollination in *L. perenne*, the following combinations of crosses and treatments were used: 1. ♀L × ♂L (before pollination stigmas were treated with pistil- or stamen-derived lectins with hemagglutination titer 1:4 from S-morphs); 2. ♀S × ♂S (before pollination stigmas were treated with pistil- or stamen-derived lectins with hemagglutination titer 1:4 from L-morphs). As a positive control the intermorph crosses without treatment was used (♀L × ♂S or ♀S × ♂L). As a negative control the self-pollination with treatment by the lectins from the same floral morph was used: before ♀S × ♂S crosses lectin from S pistils or stamens were applied, and before ♀L × ♂L crosses lectins from L pistils or stamens were used (Supplementary Fig. S1). All crossings were carried out in 16 repetitions (4 flowers each on 4 individual plants per morph). All types of crosses (both controls and all experimental lectin treatments) were made on the same four plants per morph.

## Results

### Type of self-incompatibility in L. perenne L

To determine the type of self-incompatibility in *L. perenne* three types of crosses (hand pollination) were carried out (Fig. 2): self-pollination (pollen and stigma from the same flower); intramorph crosses (pollen and stigma from the same morph, but from different individuals) and intermorph crosses (pollen and stigma from individuals of the different floral morphs).

**Fig. 2.**
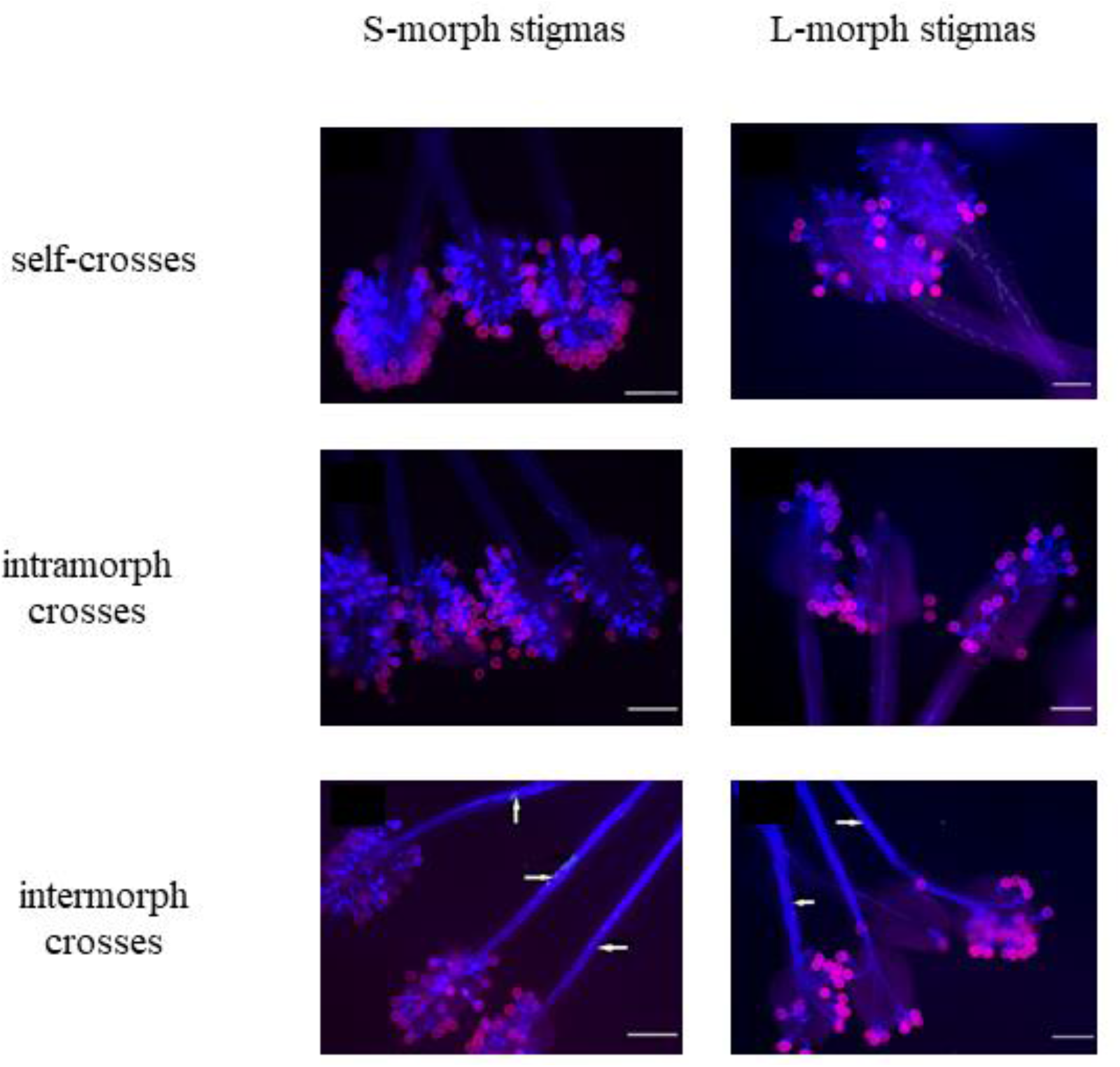
Pollen-tube growth in different types of crosses in *L. perenn*e as indicated. Pollen tubes (blue fluorescence) were visualized by aniline-blue staining; pink fluorescence indicates the pollen coat. The white arrows show pollen tubes growing in the style. Scale bars in all pictures are 200 µm

In both self-pollination and intramorph pollination (Fig. 2), the pollen tubes were arrested in the stigmas. After intermorph pollination pollen tubes grew normally in the tissues of the stigma and the style. There were no differences between using short-styled and long-styled floral morphs as mothers (Fig. 2).

### Lectin activity in different flower morphs pistils and stamens of L. perenne L

In studies on primrose (Golynskaya et al., 1976), it was found that the tissues of pistils and stamens of various flower morphs of the same species had different levels of lectin activity. In addition, for plants of the genera *Petunia* and *Eruca*, the involvement of pistil and stamen lectins in the regulation of self-incompatibility and the pollination process was assumed (Shivanna et al., 1985). In addition, it was found that lectins of some species are directly involved in the growth of the pollen tube in vitro (Matveeva et al., 2007; Sousa et al., 2013). Here we found that in both the pistils and the stamens of L-morphs and S-morphs four groups of lectins were present, which differ in carbohydrate specificity (galactose-binding and glucose-binding lectins) and in whether calcium ions are necessary for biological activity or not (Ca^2+^-dependent and Ca^2+^-independent lectins). The soluble lectin fraction contained only glucose-binding lectins, while both glucose-binding and galactose-binding lectins were found in the membrane and cellßwall fractions (Fig. 3A, C).

**Fig. 3.**
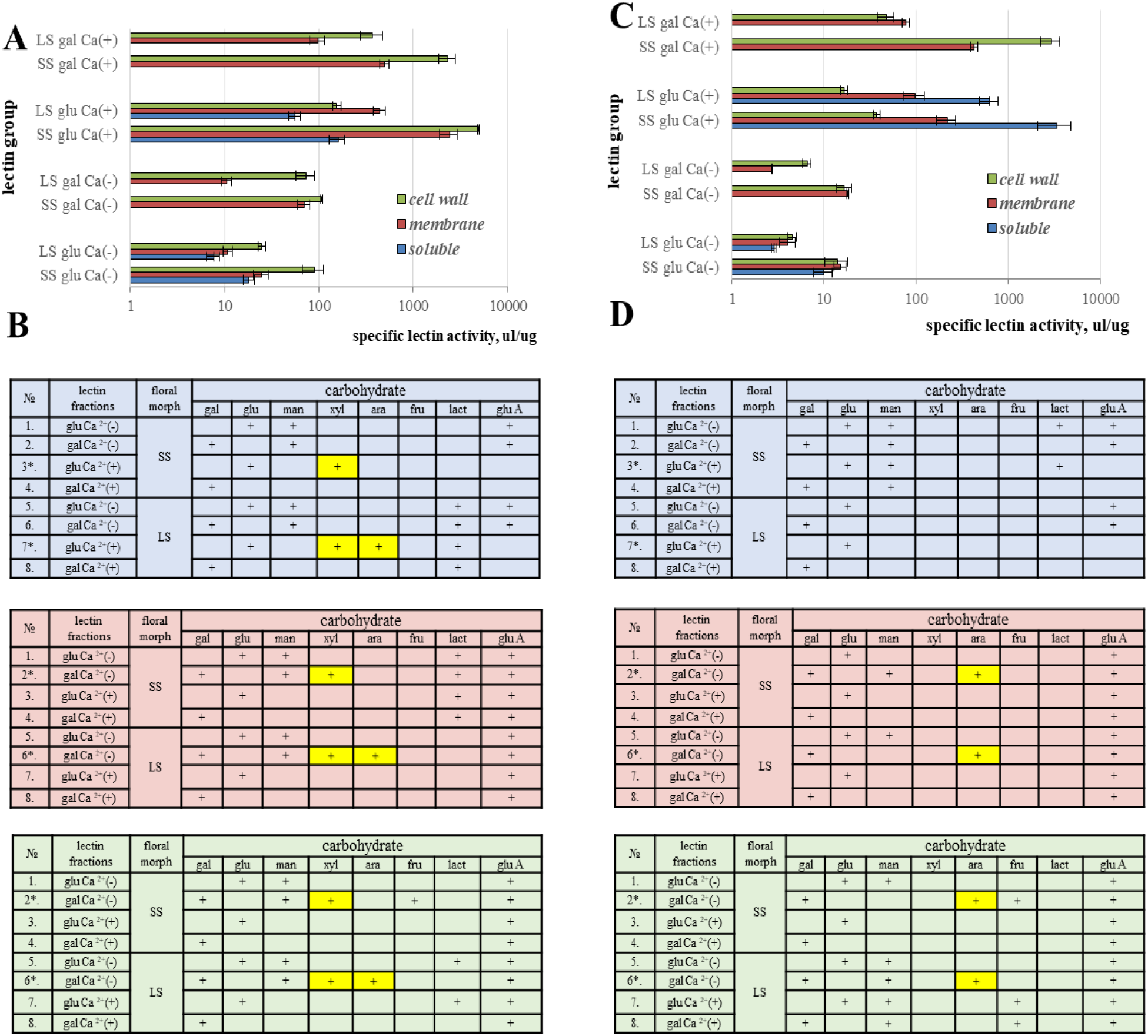
Specific lectin activity (µL/µg) and carbohydrate specificity of purified lectins extracted from pistils and stamens of different floral morphs of *L. perenne*. (A) Specific lectin activity in pistil extracts. Values are mean ± SEM of 3 biological replicates. (B) Carbohydrate specificity of pistil lectins. (C) Specific lectin activity in stamen extracts. Values are mean ± SEM of 3 biological replicates. (D) Carbohydrate specificity of stamen lectins. *S* – lectins were purified from S-morphs; *L* – lectins were purified from L-morphs; glu – glucose-binding lectins; gal – galactose-binding lectins; Ca^2+^ (+) – calcium dependent lectins; Ca^2+^ (-) – calcium independent lectins. *gal –* galactose*; glu –* glucose*; man* – mannose*, xyl –* xylose*; ara –* arabinose*; fru* – fructose*; lact –* lactose*; glu A –* glucosamine. *Blue* – carbohydrate specificity of soluble lectins; *pink* - carbohydrate specificity of membrane lectins; *green* - carbohydrate specificity of cell wall lectins. The presense of carbohydrate specificity of any sugar was show as plus (+). The purified lectin extracts which are able to overcome the self-incompatibility on pollination level are marked by asterisks (*) and yellow highlight shows the changes carbohydrate specificity of these lectins. SS: S-morph, LS: L-morph.

The amount of extracted lectins varied considerably from 0.05% to 25% of total protein depending on the lectin group or source organ. For example, the amount of Ca^2+^-independent lectins varied from 0.05% to 2% of total isolated protein, whereas the amount of Ca^2+^-dependent lectins has a wider range of variation from 0.05% to 25% of total protein content (Supplementary Table S1). However, because the biological activity of lectins depends more on their ability to bind cell-surface carbohydrates than on protein amounts, we defined total lectin activity (Supplementary Table S2) and specific lectin activity (Supplementary Table S3) as indicators of the ability to bind carbohydrates by lectins. When expressed in this way, the lectin activity is not dependent on the concentration of isolated lectins. The highest specific activity in both stamens and pistils was found for Ca^2+^-dependent glucose-specific and Ca^2+^-dependent galactose-specific lectins (Supplementary Table S3). The lectin activity in all four protein extracts from membranes was rather similar, while in cell-wall extracts stronger it depends from type of lectins: it is very high for Ca^2+^-dependent glucose-specific and Ca^2+^-dependent galactose-specific lectins and very low for another two extracts. When comparing the two floral morphs, differences in lectin activity between L-morphs and S-morphs were found (Fig. 3 A, C). In particular, for most lectin classes the extracts from pistils and stamens of S-morphs contained higher levels of specific activity than those from L-morphs.

### The carbohydrate specificity of pistil and stamen lectins extracted from different morphs of L. perenne

The biological activity of lectins can be characterized by two indicators, quantitative, i.e. lectin activity, and qualitative indicators, i.e. carbohydrate specificity. Therefore, carbohydrate specificity was determined in addition to the lectin activity. On this basis, all extracted lectins differed from each other (Fig 3B), that is they had different carbohydrate specificities. In particular, some lectin extracts (pistil-derived and stamen-derived Ca^2+^-independent galactose-binding membranes and cell wall lectins, as well as pistil-derived Ca^2+^-dependent glucose-binding soluble lectins) are able to recognize and bind xylose and/or arabinose (Fig. 3B,D). Moreover, this specificity depends on the tissue and the type of flower morph used for extraction. The corresponding lectins isolated from stamens can recognize arabinose regardless of the type of flower morph (Fig. 3D), while pistil-derived lectins of the S-morph can recognize xylose, while pistil-derived lectins of the L-morph can recognize xylose and arabinose (Fig. 3B).

### The effect of lectin treatment on self-incompatibility in heterostyled L. perenne at the pollination level

Lectins have been reported to act as signaling molecules in the pollen/stigma recognition process in self-incompatibility (Sousa et al., 2013; Sharma et al., 1985). Given the high levels of lectins with different specificities present in *L. perenne* stamens and pistils, we tested the effect of the isolated lectins on self-incompatibility in the different flower morphs of *L. perenne*. To this end, we performed self-pollination of either S-morphs or L-morphs after pretreatment of the stigmas with the different isolated lectin fractions of either the same or the other morph. Self-incompatibility was assessed by aniline-blue staining of pollinated styles. Several lectin fractions from L-plants were able to overcome the self-incompatibility in self-pollination of S-plants (Fig. 4). In particular, pre-treating the stigmas with soluble glucose-binding Ca^2+^-dependent lectins extracted from pistils, and with galactose-binding Ca^2+^-independent membrane and cell-wall lectins extracted from either pistils or stamens allowed pollen from S-plants to grow on the stigmas of their own flowers, similar to what is seen in intermorph crosses (Fig. 4). In all other cases pollen tubes were arrested in stigma (Supplementary Fig. S2).

**Fig. 4.**
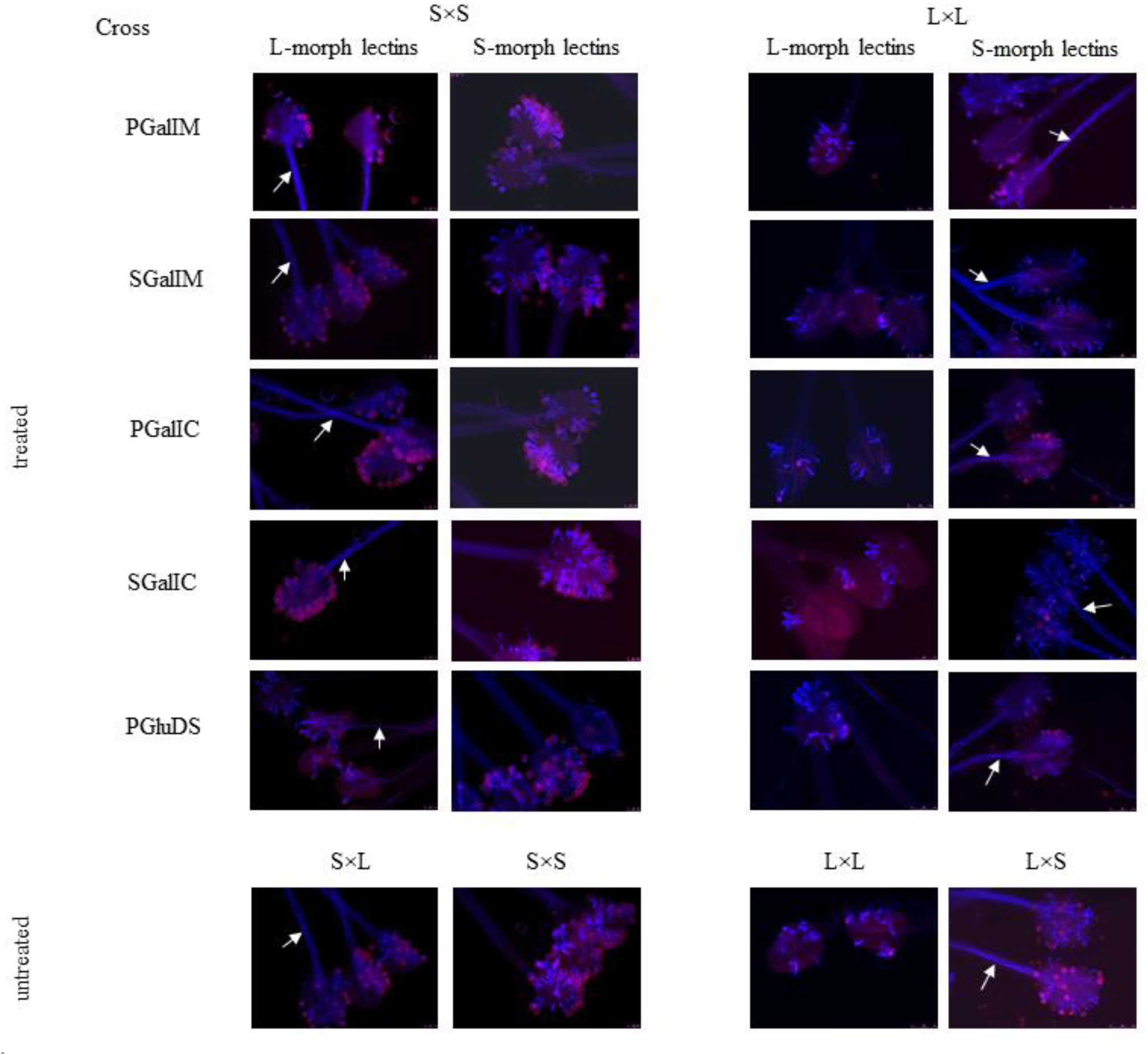
The effect of treatment by purified lectins (extracted from pistils of different floral morphs as indicated on top) on overcoming the self-incompatibility in self-crosses in heterostyled species *L. perenn*e. The pink color indicates the cover of pollen grains; the bright blue color indicates the pollen tubes. The white arrows show the pollen tubes growing in the style, when self-incompatibility has been overcome. Treatments with all other lectins that had no effect are shown in Fig. S2 and S3. S: S-morph, L: L-morph.

In self-pollination within L-morphs (L×L) similar results were observed. Also here, pre-treatment with the same lectin fractions as above (soluble glucose-binding Ca^2+^-dependent lectins extracted from pistils, and galactose-binding Ca^2+^-independent membrane and cell-wall lectins extracted from either pistils or stamens) allowed self-pollen to grow through the style (Fig. 4), while pollen tubes were arrested in all other cases (Supplementary Fig.S3). Importantly, in both morphs the pre-treatment with the relevant lectin fractions isolated from the same morph (i.e. S-stigmas pre-treated with the relevant lectin fractions from S-flowers and vice versa) did not affect the ability of pollen tubes to grow on self-stigmas (Fig. 4).

### Influence of the total lectin activity of extracts on overcoming self-incompatibility

Lectin activity is the minimum protein concentration at which agglutination occurs, i.e. complete binding of carbohydrates on the cell surface. Therefore, a more important indicator is not the concentration of the protein in the solution, but its ability to bind to carbohydrates. This indicator is called the titer or dilution. For the hemaggluination assay, a solution with a titer of 1:4 is standard used (this solution will be able to bind carbohydrates if it is diluted 4 times) because the erythrocyte suspension dilutes the initial lectin solution.

To test the effectiveness of the different lectins on overcoming self-incompatibility, we repeated the above experiment using dilution series of the different lectins. We used the three pistil-derived lectin fractions that enabled successful self-fertilization (PGluDS, PGalIC and PGalIM) before, and treated stigmas with increasing dilutions of these (1:4, 1:8, 1:16 and 1:32), before self-pollinating them. In both morphs, PGluDS lectins are only able to overcome self-incompatibility at high titer (1:4 and partially 1:8) (Fig. 5) By contrast, PGalIM and PGalIC lectins overcame self-incompatibility even at a titer of 1:16 and partially at a titer of 1:32 (Fig. 5). Thus, membrane and cell wall-derived lectins are more effective than soluble lectins.

**Fig. 5.**
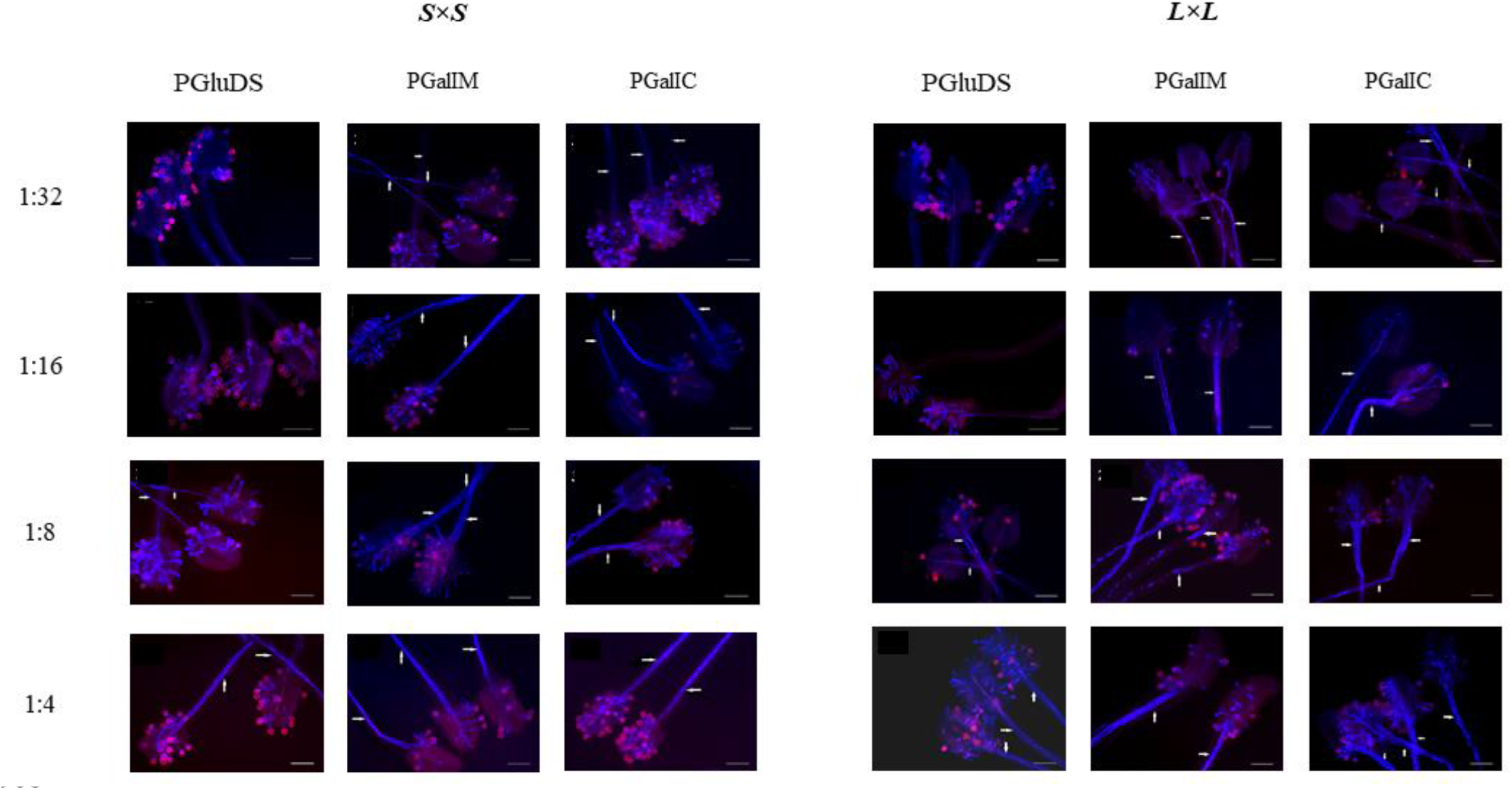
Relative activity of different pistillate lectin fractions in overcoming self-incompatibility in self-crosses of *L. perenne* as assessed by a dilution series. Dilution series based on hemagglutination titer of the following lectins purified from pistils of the other morph were used : *PGluDS* - soluble glucose-binding Ca^2+^-dependent lectin, *PGalIM* – membrane galactose-binding Ca^2+^-independent lectins and *PGalIC –* cell wall galactose-binding Ca^2+^-independent lectins. The stigmas were treated with lectin extracts, diluted to a titer of 1:4, 1:8, 1:16 or 1:32 as indicated on the left, before self-pollination. The pink color indicates the cover of pollen grains; the bright blue color indicates the pollen tubes. The white arrows show the pollen tubes growing in the style. Scale bars in all pictures are 200 µm. S: S-morph, L: L-morph.

### The effect of lectins on self-incompatibility in heterostyled L. perenne at the fertilization level

To test whether the observed effect of lectin treatment on self-incompatibility at the level of pollen-tube growth could be translated into successful fertilization of ovules, we repeated the above experiment with the five lectin extracts that could overcome self-incompatibility at the pollen-tube level and measured capsule formation and seed set. To minimize the influence of genetic variation between the mother plants, we performed all crosses in a multiply paired design. For this, four S and four L plants were chosen, and four flowers each per plant were subjected to each of the tested treatments and controls, ensuring that different treatment effects indeed reflected the different activities of the various lectin fractions.

In both morphs, pre-treating stigmas with either of three lectin fractions derived from pistils of the other morph could overcome self-incompatibility and lead to seed formation after self-pollination (Table 1, Fig. 6). By contrast, the corresponding stamen-derived lectins or the pistil-derived lectins from the same morph had no effect. The active fractions were pistil-derived soluble glucose-binding calcium-dependent lectins (PGluDS), pistil-derived galactose-binding Ca^2+^-independent cell wall lectins (PGalIC), and pistil-derived galactose-binding calcium-independent membrane lectins (PGalIM). Of these, PGalIC had the strongest effect in both morphs. In particular, after PGalIC treatment 14 of the 16 tested S flowers formed capsules with an average of 7 seeds each, while 12 of the 16 tested L flowers formed capsules with an average of 7 seeds each. By contrast, PGalIM treatment with the weakest effect led only to the formation of 10 capsules from 16 treated S flowers with on average 4 seeds, and to the formation of 8 capsules from the 16 treated L flowers with on average 4 seeds. Thus, these three lectin fractions can not only overcome the self-incompatibility barrier at the level of the stigma, but also lead to successful self-fertilization. By contrast, the stamen-derived lectin fractions that could overcome self-incompatibility at the stigma level (galactose-binding Ca^2+^-independent membrane lectins [SGalIM] and galactose-binding Ca^2+^-independent cell wall lectins [SGalIC]) failed to enable self-fertilization. At the same time, even the strongest lectin effect fell short of the capsule formation and seed set obtained in intermorph crosses (16 out of 16 capsules formed, with 8 seeds on average in both morphs), though not by much in S plants. In addition, we determined the viability of the obtained seeds at the stage of seedlings and adult plants. The best viability was obtained after treatment by PGalIC lectins, reaching almost the level of the intermorph-cross positive control, with 85.3% survival at the seedling and 67.6% at the adult stage compared with the intermorph-cross control values of 87.5% and 77.5%, respectively. By contrast, after treatment by PGalIM lectins survival rates were only 21.7% and 17.4% at the seedling and adult-plant stages, respectively.

**Table 1.** The effect of lectin treatment in overcoming the self-incompatibility in heterostyled *L. perenne* at the level of seed set.

| Type of crosses | Experiment |  |  |  |  |  |  |  |  |  |  |
| --- | --- | --- | --- | --- | --- | --- | --- | --- | --- | --- | --- |
|  | positive control | Type of lectin extract |  |  |  |  |  |  |  |  |  |
|  |  | PGalIM |  | SGalIM |  | PGalIC |  | SGalIC |  | PGluDS |  |
|  |  | morph origin |  |  |  |  |  |  |  |  |  |
|  |  | S | L | S | L | S | L | S | L | S | L |
| S×S |  |  |  |  |  |  |  |  |  |  |  |
| the fruit set | 16 | - | 10 | - | - | - | 14 | - | - | - | 12 |
| the average seed set in each capsule | 8.5±0.67 | - | 4.2±1.55 | - | - | - | 6.8±0.42 | - | - | - | 5.55±0.85 |
| the total seed set | 126 | - | 45 | - | - | - | 98 | - | - | - | 70 |
| surviving seedlings (% of total seed set) | 111 (88.6) | - | 31 (68.9) | - | - | - | 84 (85.6) | - | - | - | 14 (20.2) |
| surviving mature plants (% of total seed set) | 96 (76.2) | - | 23 (51.1) | - | - | - | 67 (68.4) | - | - | - | 11 (15.7) |
| L×L |  |  |  |  |  |  |  |  |  |  |  |
| the fruit set | 16 | 8 | - | - | - | 12 | - | - | - | 10 | - |
| the average seed set in each capsule | 8.6±0.72 | 4.0±1.64 | - | - | - | 6.5±0.51 | - | - | - | 5.75±0.55 | - |
| the total seed set | 130 | 33 | - | - | - | 72 | - | - | - | 58 | - |
| surviving seedlings (% of total seed set) | 114 (87.7) | 22 (66.7) | - | - | - | 61 (84.7) | - | - | - | 12 (20.7) | - |
| surviving mature plants (% of total seed set) | 101 (77.7) | 17 (51.5) | - | - | - | 49 (68.0) | - | - | - | 10 (17.4) | - |

**Note:**
The crosses were carried out in 2015-2016, with 16 biological replicates per cross type. Crosses were performed in a paired manner, i.e. four flowers each on the same four plants were treated with either S- or L-derived lectins before pollination, in order to exclude variation between mother plants as the cause for the different effects of S- and L-derived lectins. Thus, all of the treatments shown were performed on the same four mother plants per morph. Only lectin extracts that had a positive effect at the level of pollen-tube growth were used. S: S-morph, L: L-morph.
As a negative control the self-pollination with treatment by the lectins from the same floral morph was used: before ♀S × ♂S crosses lectin from SS pistils or stamens were applied, and before ♀L × ♂L crosses lectins from L pistils or stamens were used. These treatments resulted in no seed set.
PGalIM – pistillate galactose-binding Ca<sup>2+</sup>-independent membrane lectins; SGalIM – staminate galactose-binding Ca<sup>2+</sup>-independent membrane lectins; PGalIC – pistillate galactose-binding Ca<sup>2+</sup>-independent cell wall lectins; SGalIC – staminate galactose-binding Ca<sup>2+</sup>-independent cell wall lectins; PGluDS – pistillate glucose-binding Ca<sup>2+</sup>-dependent soluble lectins.

**Fig. 6.**
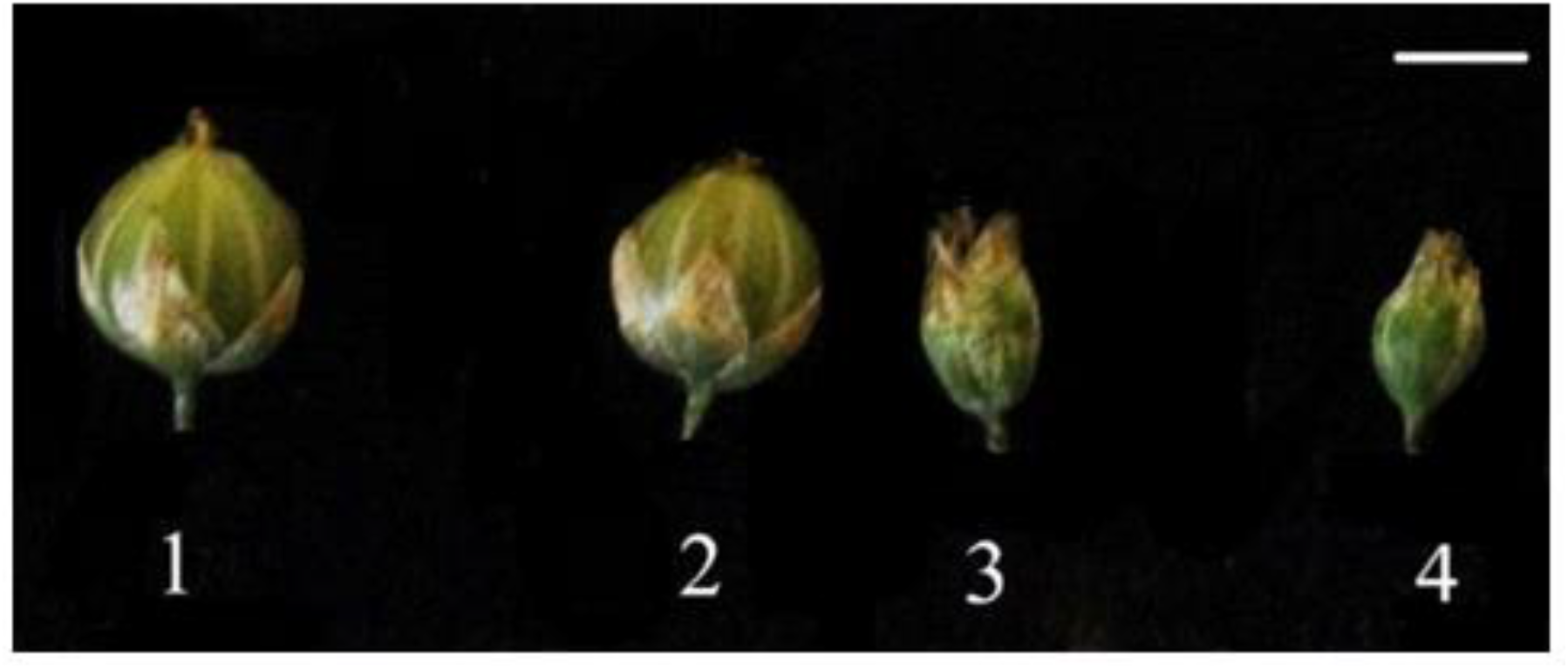
Capsule formation after overcoming self-incompatibility by lectin treatment. **1**– positive control (S×L); **2,3** – capsules formed after treatment of S-morph stigmas by pistil-derived (2) or stamen-derived (3) galactose-binding calcium-independent lectins (PGalIC) from L flowers, followed by self-pollination; **4** –negative control (S×S crosses after treatment with pistillate PGalIC lectins extracted from S flowers). Scale bar is 5 mm.

### Prolonged effect of lectins in overcoming self-incompatibility in L. perenne

We sought to determine whether the effect of lectin treatment on self-incompatibility was limited only to the treated generation of plants, or whether it might also affect the offspring of the treated plants. To this end, self-pollination of the obtained plants from lectin-treated selfed S-morphs was carried out over three generations.

When F_1_ progeny obtained after selfing short-styled plants treated with PGluDS were self-pollinated again without any further lectin treatment, 5 out of 16 self-pollinated F_1_ flowers formed capsules, with an average of 5 seeds across the 5 capsules, yet none of the seeds was viable (Table 2). Thus, there was no transgenerational effect of PGluDS treatment. By contrast, the F_1_ progeny of selfed PGalIM lectin-treated plants retained higher self-compatibility, forming 8 out of 16 capsules with an average seed set of 5 seeds per capsule. F_2_ plants derived from these still retained partial self-compatibility (4 out of 16 capsules formed, 4 seeds on average), yet none of the seeds were viable. PGalIC treatment had the longest-lasting effect. High levels of self-compatibility were observed in the F_1_ and F_2_ plants derived from selfed PGalIC-treated plants, and even F_3_ plants still retained self-compatibility, with 6 out of 16 capsules formed, containing 4 seeds each on average, and these seeds gave rise to a high proportion of viable F_4_ plants. Thus, treatment with PGalIC lectins causes a long-lasting suppression of self-incompatibility that is transmitted to at least three untreated generations, indicating as a prolonged effect of lectin treatment.

**Table 2.**
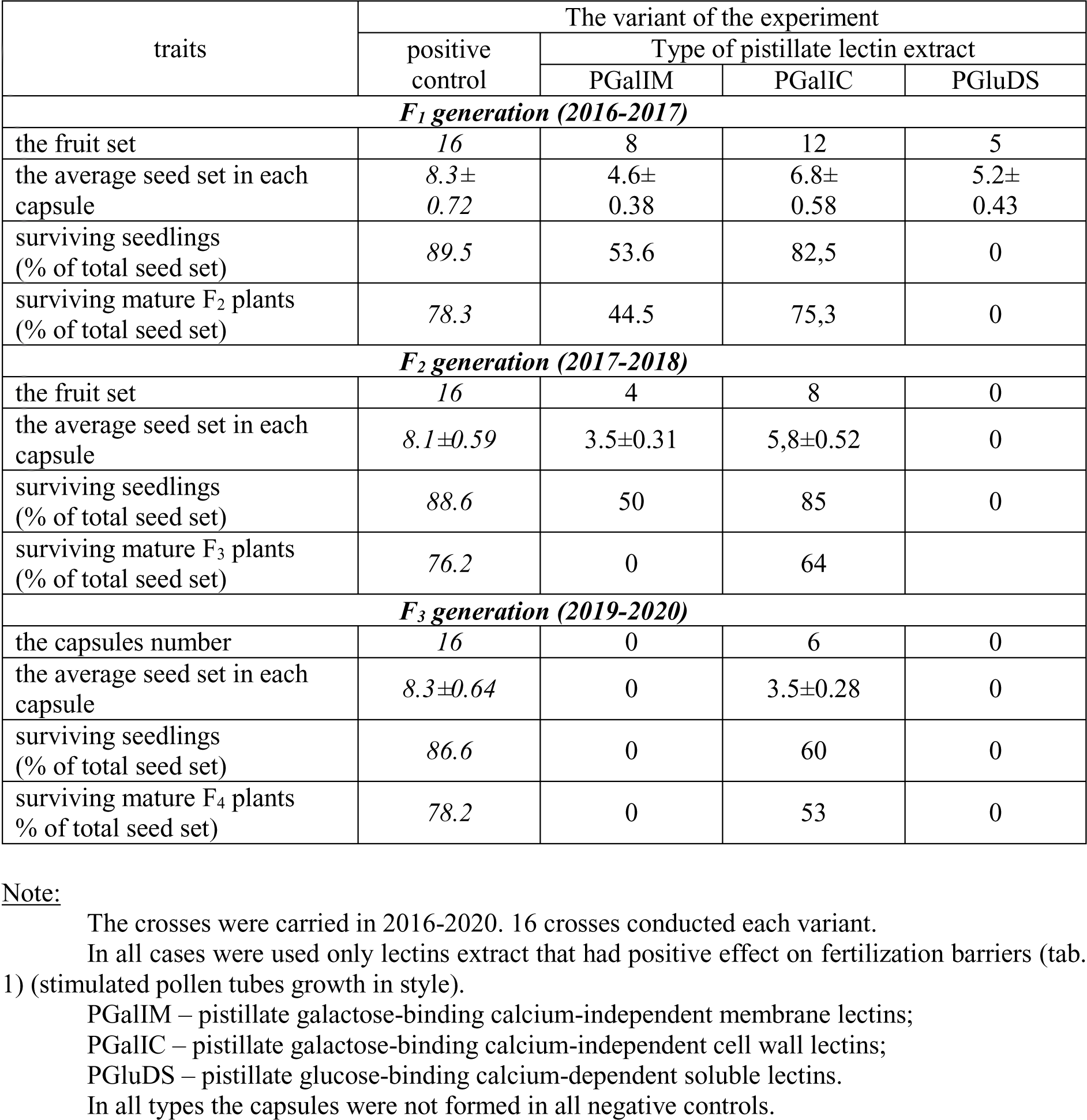
The prolonged effect of lectin treatment on self-incompatibility in S-morph of *L. perenne* L.

## Discussion

In this work we show that lectins with different cellular localization – soluble, membrane and cell wall lectins – are present in pistil and stamen tissues of *L. perenne* flowers. Each of these fractions (except soluble lectins) contained at least four different lectin types with different carbohydrate specificities (glucose-binding and galactose-binding lectins) and the different requirements for calcium ions for their carbohydrate-binding activity (Ca^2+^-dependent and Ca^2+^-independent lectins). There was variation in the specific activity of lectins isolated from different floral morphs. Also, for pistil-derived lectins the carbohydrate specificity depended on the type of floral morph they were isolated from. The organ- and morph-specific distribution of these proteins suggests that they could contribute to the self-incompatibility mechanism. To test this, stigmas were treated with the different lectin fractions before self- and intra-morph pollination, and self-incompatibility was assessed both at the level of pollen-tube growth and of capsule formation and seed set. Indeed, all galactose-binding Ca^2+^-independent lectins extracted from membranes and cell walls of either pistils or stamens, as well as soluble glucose-binding Ca^2+^-dependent lectins from pistils are capable of overcoming the self-incompatibility reaction at the level of pollen-tube growth, allowing robust pollen-tube growth through the styles after self- and intra-morph pollination in both floral morphs of *L. perenne*. However, seeds were only formed after treatment with the corresponding pistil-derived lectins isolated from the opposite morph; neither stamen-derived lectins nor pistil-derived lectins from the same morph had any effect on allowing self-fertilization. This morph-specific activity is particularly important as it argues against a mere general germination- or growth-stimulating effect of the respective lectin fractions and supports the conclusion that the isolated lectins can indeed specifically modulate the heteromorphic self-incompatibility mechanism in *L. perenne*. This activity is most likely due to the carbohydrate specificity of the lectins for the following reasons. First, only the five lectins that can promote self-pollen tube germination and growth can recognize and bind arabinose and/or xylose. Second, all pistil-derived lectins that enable self-fertilization can bind xylose, while the stamen-derived lectins cannot. Finally, all lectins from cell wall and membranes including PGalIM and PGalIC that have a positive effect on overcoming SI can bind glucosamine. This suggests that these proteins have at least one lectin domain from Nictaba or Jacalin Related Lectin families because only these two lectins groups can bind these specific carbohydrates (De Coninck and Van Damme. 2021).

Surprisingly, the pistil-derived lectin fractions discussed above could not only overcome self-incompatibility in the treated generation, but for two lectins, this effect persisted over up to three untreated generations. This strongly suggests that the lectin treatment possibly can stimulated epigenetic changes. A possible epigenetic effect of lectins could be based on the ability of some lectins to bind histones, as has been demonstrated for Nictaba lectins from tobacco (Schouppe et al., 2011). Thus, treatment with PGalIM and PGalIC lectins may affect the expression of incompatibility genes leading to a transgenerational effect that allows self-fertilization of subsequent untreated generations.

## Conclusion

This study shows that pistil-derived galactose-specific Ca^2+^-independent lectins can overcome self-incompatibility in heterostylous *L. perenne*. This may be due to the presence of some lectins that are specific for arabinose and xylose. Our results suggest that lectins can participate in signaling pathways for recognition at the pollen-stigma interface and are involved in the regulation of the self-incompatibility process. However, it remains to be determined which lectins are involved in heteromorphic self-incompatibility in *L. perenne*.

