## Supplemental Figure 1, Supplemental Table 1, Supplemental Table 2, Supplemental Table 3, Supplemental Figure 2, Supplemental Figure 3 for "Lectins in pistils play a key role in self-incompatibility in the heterostylous *Linum perenne*"

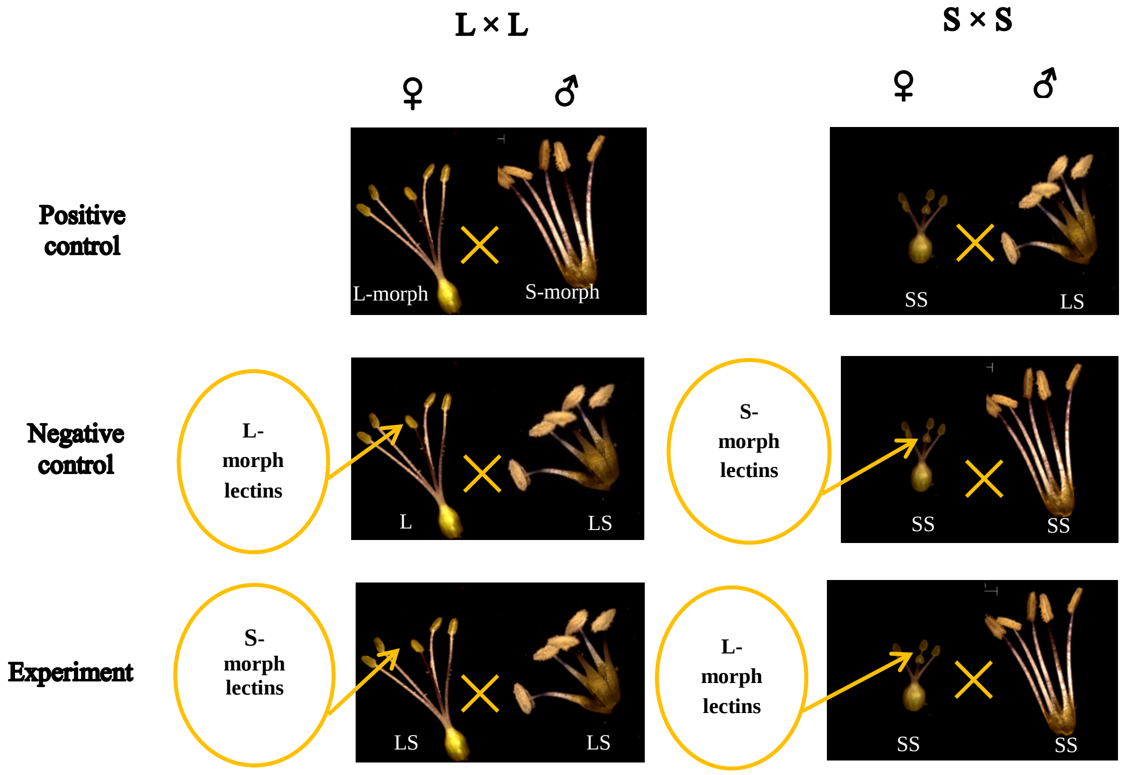


**Fig. S1.** The scheme of crosses. Two types of crosses were conducted – S×S and L×L. In all crosses were experiment and two types of controls: positive and negative controls. In S×S crosses negative controls were crosses after treatment by lectins extracted from S-morph and positive control in these crosses were S×L crosses. In experiment stigmas before pollinated were treated by lectins extracted from L-morphs. In L×L crosses negative controls were crosses after treatment by lectins extracted from L-morph and positive control in these crosses were L×S crosses. In experiment stigmas before pollinated were treated by lectins extracted from S-morphs. In all cases were used lectins extracted from pistils and stamens

***Table S1.***

Total protein concentration (mg/ml) at the different stages of lectin extraction and purification from pistils and stamens of different *L. perenne* floral morphs

| extracts | Soluble lectins | | | | Membrane lectins | | | | Cell wall lectins | | | |
|---|---|---|---|---|---|---|---|---|---|---|---|---|
| | ♀ S | ♂ S | ♀ L | ♂ L | ♀ S | ♂ S | ♀ S | ♂ S | ♀ S | ♂ S | ♀ S | ♂ S |
| **After extraction (total protein amount)** | **68.52±4.34** | **56.73±2.55** | **63.61±5.33** | **55.16±3.87** | **9.97±0.51** | **13.07±2.02** | **10.80±0.72** | **13.44±1.44** | **10.94±1.27** | **11.47±2.25** | **7.24±1.50** | **6.55±0.75** |
| **After extraction and salting out of proteins, after dialysis** | 21.65±1.09 | 21.18±2.77 | 28.21±4.74 | 26.04±1.48 | 4.74±0.69 | 5.39±0.55 | 6.26±0.88 | 7.13±0.75 | 5.61±0.43 | 6.58±0.89 | 4.61±0.79 | 3.55±063 |
| **After lectin binding to the glucose-specific resin** | 4.22±0.34 | 4.29±0.57 | 1.90±0.47 | 3.14±0.56 | 1.74±0.16 | 1.17±0.28 | 1.13±0.38 | 0.99±0.15 | 1.11±0.26 | 0.58±0.10 | 1.54±0.12 | 0.74±0.10 |
| **After lectin binding to the galactose-specific resin** | 0.23±0.08 | 0.72±0.14 | 0.46±0.07 | 0.84±0.11 | 0.69±0.13 | 0.62±0.11 | 0.73±0.07 | 0.49±0.12 | 0.77±0.10 | 0.67±0.13 | 0.36±0.10 | 0.40±0.08 |
| **After lectin elution from the resin by glucose and dialysis** | 3.65±0.43 | 3.65±0.85 | 4.41±0.65 | 5.61±0.32 | 2.78±0.43 | 1.63±0.28 | 3.04±0.71 | 2,13±0.43 | 3.33±0.85 | 5.17±1.09 | 5.28±0.54 | 7.24±0.66 |
| **After lectin elution from the resin by galactose and dialysis** | - | - | - | - | ***1.91±0.24**** | ***1.59±0.79**** | ***6.26±0.69**** | ***5.58±0.32**** | ***4.84±0.11**** | ***8.22±1.27**** | ***3.87±0.75**** | ***10.10±1.32**** |
| **After lectin elution from the glucose-specific resin by EDTA and dialysis** | ***0.43±0.07**** | 0.39±0.10 | ***0.61±0.09**** | 0.48±0.14 | 0.47±0.09 | 0.65±0.12 | 0.61±0.09 | 0.74±0.10 | 0.59±0.13 | 0.87±0.07 | 0.83±0.16 | 0.99±0.17 |
| **After lectin elution from the galactose-specific resin by EDTA and dialysis** | - | - | - | - | 1.05±0.11 | 0.62±0.06 | 1.25±-0.07 | 0.85±0.09 | 0.94±0.22 | 0.20±0.03 | 1.65±0.48 | 1.59±0.06 |
| **After washed the resin from EDTA** | - | - | - | - | - | - | - | - | - | - | - | - |

**Note:**

***White*** – total extracts;
***Yellow*** – remaining total protein in extracts after lectin binding to resins;
***Blue*** – purified lectins eluted from resins by carbohydrates;
***Pink*** – purified lectins eluted from resins by EDTA;

♀ - extract from pistils;
♂ - extract from stamens
***SS*** - short-styled floral morphs;
***LS*** - long-styled floral morphs;

***The purified lectin extracts which are able to overcome the self-incompatibility on pollination level are shown in bold italics and marked by asterisks (*).***

***Table S2.***

Total lectin activity (titer) at the different stages of lectin extraction and purification from pistils and stamens of different *L. perenne* floral morphs

| **extracts** | **Soluble lectins** | | | | **Membrane lectins** | | | | **Cell wall lectins** | | | |
|---|---|---|---|---|---|---|---|---|---|---|---|---|
| | **♀S** | **♂S** | **♀L** | **♂L** | **♀S** | **♂S** | **♀S** | **♂S** | **♀S** | **♂S** | **♀S** | **♂S** |
| **After extraction (total protein amount)** | **256** | **64** | **128** | **16** | **256** | **64** | **64** | **16** | **8192** | **4096** | **2048** | **1024** |
| **After extraction and salting out of proteins, after dialysis** | 64 | 32 | 32 | 8 | 128 | 32 | 32 | 8 | 256 | 64 | 128 | 32 |
| **After lectin binding to the glucose-specific resin** | - | - | - | - | 16 | 8 | 8 | 4 | 16 | 8 | 8 | 4 |
| **After lectin binding to the galactose-specific resin** | - | - | - | - | - | - | - | - | - | - | - | - |
| **After lectin elution from the resin by glucose and dialysis** | 64 | 32 | 32 | 16 | 64 | 16 | 32 | 8 | 256 | 64 | 128 | 32 |
| **After lectin elution from the resin by galactose and dialysis** | - | - | - | - | ***128**** | ***32**** | ***64**** | ***16**** | ***512**** | ***128**** | ***256**** | ***64**** |
| **After lectin elution from the glucose-specific resin by EDTA and dialysis** | ***64**** | 1024 | ***32**** | 256 | 1024 | 128 | 256 | 64 | 256 | 32 | 128 | 16 |
| **After lectin elution from the galactose-specific resin by EDTA and dialysis** | - | - | - | - | 512 | 256 | 128 | 64 | 2048 | 512 | 512 | 64 |
| **After washed the resin from EDTA** | - | - | - | - | - | - | - | - | - | - | - | - |

**Note:**

***White*** – total extracts;
***Yellow*** – remaining total protein in extracts after lectin binding to resins;
***Blue*** – purified lectins eluted from resins by carbohydrates;
***Pink*** – purified lectins eluted from resins by EDTA;

♀ - extract from pistils;
♂ - extract from stamens
***SS*** - short-styled floral morphs;
***LS*** - long-styled floral morphs;

***The purified lectin extracts that are able to overcome the self-incompatibility on pollination level are shown in bold italics and marked by asterisks (*).***

***Table S3.***

Specific lectin activity (μl/μg) at the different stages of lectin extraction and purification
from pistils and stamens of different *L. perenne* floral morphs

| **extracts** | **Soluble lectins** | | | | **Membrane lectins** | | | | **Cell wall lectins** | | | |
|---|---|---|---|---|---|---|---|---|---|---|---|---|
| | ♀ **S** | ♂ **S** | ♀ **L** | ♂ **L** | ♀ **S** | ♂ **S** | ♀ **S** | ♂ **S** | ♀ **S** | ♂ **S** | ♀ **S** | ♂ **S** |
| **After extraction (total protein amount)** | **3.74±0.04** | **1.13±0.04** | **2.01±0.07** | **0.29±0.03** | **25.81±1.33** | **5.12±0.66** | **5.96±0.37** | **1.19±0.02** | **748.66± 8.73** | **359.12± 18.24** | **307.90± 55.46** | **160.99± 18.42** |
| **After extraction and salting out of proteins, after dialysis** | 2.95±0.05 | 1.51±0.02 | 1.20±0.17 | 0.31±0.03 | 28.33±4.07 | 6.08±0.62 | 5.36±0.82 | 1.15±0.12 | 46.23±3.41 | 10.08±1.21 | 27.88±1.54 | 9.66±1.70 |
| **After lectin binding to the glucose-specific resin** | - | - | - | - | 10.27±1.93 | 11.11±2.42 | 7.59±1.45 | 4.24±0.56 | 16.22±3.43 | 14.61±2.60 | 6.56±1.95 | 5.52±0.99 |
| **After lectin binding to the galactose-specific resin** | - | - | - | - | - | - | - | - | - | - | - | - |
| **After lectin elution from the resin by glucose and dialysis** | 18.15±2.3 8 | 9.89±2.24 | 7.63±1.18 | 2.87±0.17 | 24.51±4.44 | 15.02±2.33 | 10.83±1.25 | 4.10±0.80 | 89.58±23.01 | 14.16±3.89 | 24.77±2.34 | 4.51±0.45 |
| **After lectin elution from the resin by galactose and dialysis** | - | - | - | - | ***69.68±9.89**** | ***18.44±0.55**** | ***10.52±1.22**** | ***2.71*±0.06*** | ***105.85± 2.32**** | ***16.64±3.07**** | ***73.26±16.78**** | ***6.54±0.72**** |
| **After lectin elution from the glucose-specific resin by EDTA and dialysis** | ***159.17± 30.20**** | 3371.90± 1314.65 | ***55.19± 8.06**** | 630.07± 144.44 | 2417.51± 516.50 | 217.86± 51.42 | 441.52± 64.46 | 97.69±24.99 | 486.69± 102.32 | 37.72±3.06 | 154.49±15.81 | 16.69±1.58 |
| **After lectin elution from the galactose-specific resin by EDTA and dialysis** | - | - | - | - | 498.26± 53.43 | 421.22± 39.88 | 96.96±18.08 | 77.22±7.36 | 2301.31± 452.99 | 2902.37± 707.86 | 372.01± 99.33 | 47.41±9.35 |
| **After washed the resin from EDTA** | - | - | - | - | - | - | - | - | - | - | - | - |

**Note:**

***White*** – total extracts;
***Yellow*** – remaining total protein in extracts after lectin binding to resins;
***Blue*** – purified lectins eluted from resins by carbohydrates;
***Pink*** – purified lectins eluted from resins by EDTA;

♀ **-** extract from pistils;
♂ - extract from stamens
***S*** - short-styled floral morphs;
***L*** - long-styled floral morphs;

***The purified lectin extracts that are able to overcome the self-incompatibility on pollination level are shown in bold italics and marked by asterisks (*).***

Ca$^{2+}$-independent lectins | Ca$^{2+}$-dependent lectins

pistillate | staminate | pistillate | staminate

***glucose-binding lectins***

*soluble lectins*

*membrane lectins*

*cell wall lectins*

***galactose-binding lectins***

*membrane lectins*

*cell wall lectins*

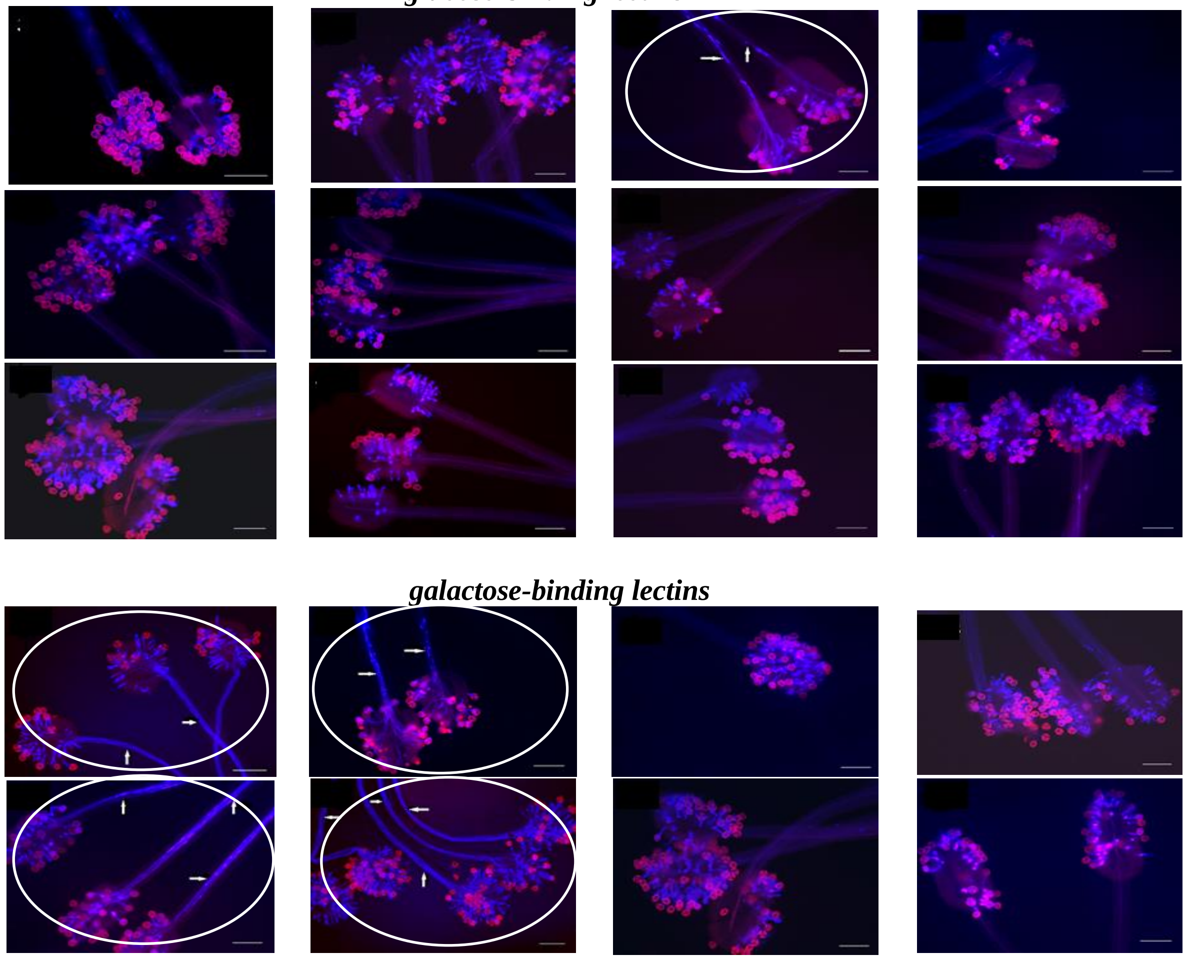

**Fig. S2.** The effect of treatment by purified lectins to overcome the self-incompatibility in **S×S** self-crosses in *L. perenne* L. The pink color indicates the cover of pollen grains; the bright blue color indicates the pollen tubes. The white arrows show the pollen tubes growth in style (the overcoming of self-incompatibility). Scale bars in all pictures are 200 µm.

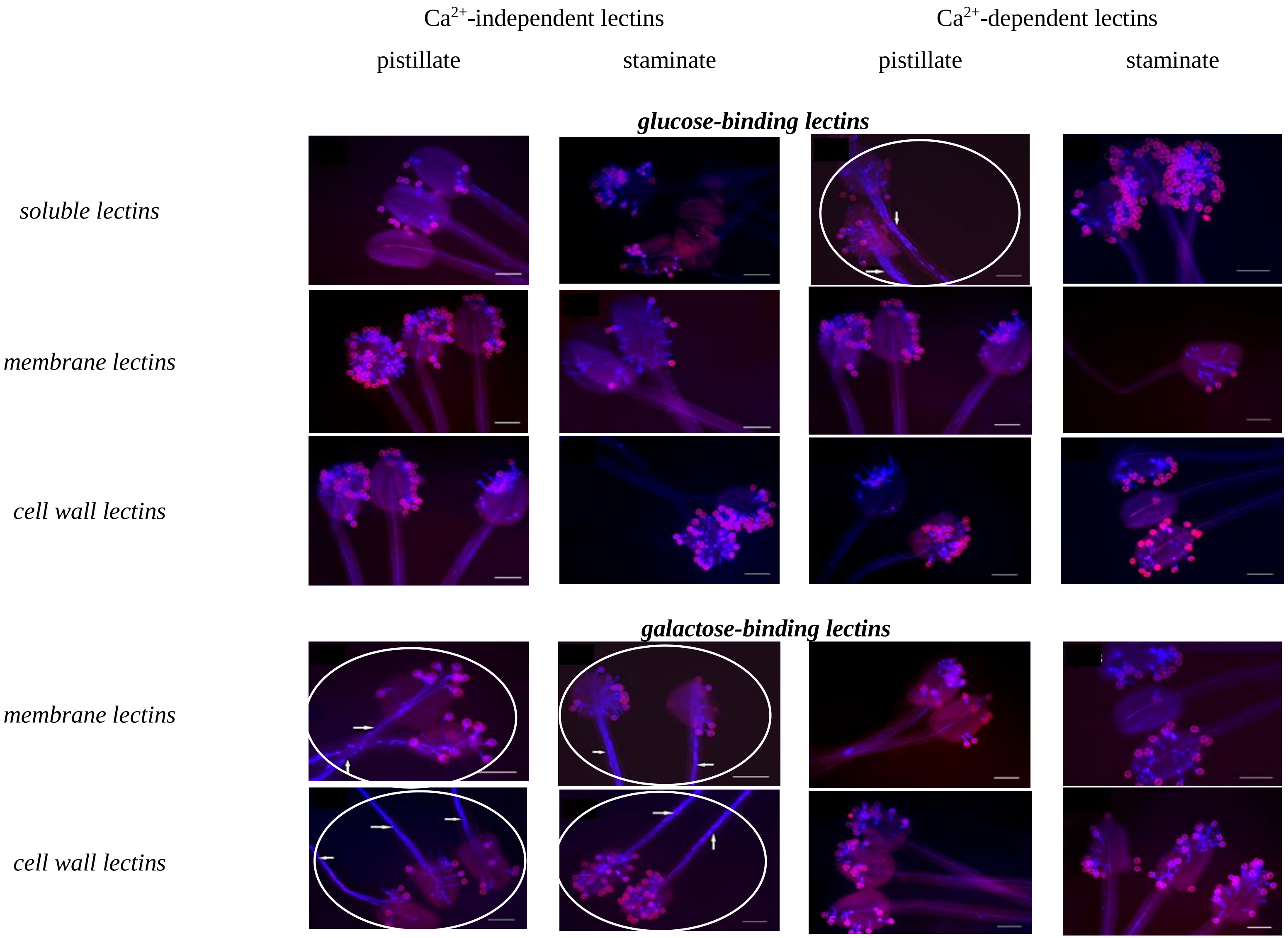


**Fig. S3.** The effect of treatment by purified lectins to overcome the self-incompatibility in **L×L** self-crosses in *L. perenne* L. The pink color indicates the cover of pollen grains; the bright blue color indicates the pollen tubes. The white arrows show the pollen tubes growth in style (the overcoming of self-incompatibility). Scale bars in all pictures are 200 µm.
